# Stepwise evolution and exceptional conservation of ORF1a/b overlap in coronaviruses

**DOI:** 10.1101/2021.06.14.448413

**Authors:** Han Mei, Anton Nekrutenko

## Abstract

The programmed frameshift element (PFE) rerouting translation from *ORF1a* to *ORF1b* is essential for propagation of coronaviruses. A combination of genomic features that make up PFE—the overlap between the two reading frames, a slippery sequence, as well as an ensemble of complex secondary structure elements—puts severe constraints on this region as most possible nucleotide substitution may disrupt one or more of these elements. The vast amount of SARS-CoV-2 sequencing data generated within the past year provides an opportunity to assess evolutionary dynamics of PFE in great detail. Here we performed a comparative analysis of all available coronaviral genomic data available to date. We show that the overlap between *ORF1a* and *b* evolved as a set of discrete 7, 16, 22, 25, and 31 nucleotide stretches with a well defined phylogenetic specificity. We further examined sequencing data from over 350,000 complete genomes and 55,000 raw read datasets to demonstrate exceptional conservation of the PFE region.

---

Coronaviruses have large 26-32 kbp +-strand RNA genomes. The initial ⅔ of the genome is occupied by an open reading frame (ORF) *ORF1ab* encoding nsps essential for coronaviral life cycle. As the designation “ab” suggests it contains two reading frames with the 3’-end of *ORF1a* overlapping with the 5’-terminus of *ORF1b. ORF1b* is in −1 phase relative to ORF1a and translated via the −1 programmed ribosomal frameshifting controlled by the programmed frameshift element (PFE). As *ORF1b* encodes crucial components of coronavirus transcription/replication machinery including the RNA-dependent RNA polymerase (RdRp) disrupting PFE abolishes viral replication completely (Brierley 1995; Plant et al. 2010; Sola et al. 2015; Kelly et al. 2020). PFE consists of three consecutive elements: (1) an attenuator loop, (2) the “NNN WWW H” slippery heptamer, and (3) a pseudoknot structure (Kelly et al. 2020; Huston et al. 2021). The sequence and structural conformation of these elements determines the efficiency of the frameshift event, which ranges from 15 to 30% in SARS-CoV and SARS-CoV-2 (Baranov et al. 2005; Kelly et al. 2020). Because disruption of FSE arrests viral replication it is a promising therapeutic target. As a result a number of recent studies have scrutinized its characteristics (reviewed in (Rangan et al. 2021)) revealing a fluid secondary structure (Iserman et al. 2020; Ziv et al. 2020; Huston et al. 2021). In addition to secondary structures FSE harbors the overlap between *ORF1a* and *ORF1b*. It is defined as the stretch of sequence from “H” in the slippery heptamer to the stop codon of *ORF1a*. The position of *ORF1a* stop codon determines overlap length. For example, in SARS-CoV-2 it is 16 bp while in mouse hepatitis virus (MHV) it is 23 nt (Plant et al. 2010).

Our group has been interested in the evolutionary dynamics of overlapping coding regions (Nekrutenko et al. 2005; Chung et al. 2007; Szklarczyk et al. 2007). The vast amount of newly generated sequence and functional data—a result of the current SARS-CoV-2/COVID-19 pandemic—provides an opportunity to reexamine our current knowledge. The length of the *ORF1a* and *ORF1b* overlap is phylogenetically conserved. It evolved in a stepwise manner, where the changes in the overlap length are results of the loss of *ORF1a* stop codons leading to *ORF1a* extension, and the acquisition of insertions and deletions causing early stops of ORF1a.

Distance-based methods had shown that the δ-coronavirus genus was an early split-off lineage compared to α-, β-, and γ-coronavirus (Fig. 1). Comparisons of the RdRp, 3CL^pro^, HEL, M, and N proteins suggested that γ-was more closely related to δ-coronavirus, while α- and β-coronavirus cluster together forming a distant clade (de Groot et al. 2012; Lau et al. 2012; Woo et al. 2012; Coronaviridae Study Group of the International Committee on Taxonomy of Viruses 2020). However, comparing the S protein trees, α- and δ-coronavirus share a higher amino acid identity, while β- and γ-coronavirus cluster together (Lau et al. 2012). Due to this we initially assumed that α, β, and γ form an unresolved trifurcation (Fig. 1). To assess all possible configurations within this region we surveyed all genomic sequences of family Coronaviridae available from the National Center for Biotechnology Information (NCBI; see Methods). The distribution of overlap lengths among 4,904 coronaviral genomes (Table S1) is shown in Fig. 2. There are five distinct overlap length groups (7, 16, 22, 25, and 31 nt) with clear taxonomic specificity.

**Fig. 1.**
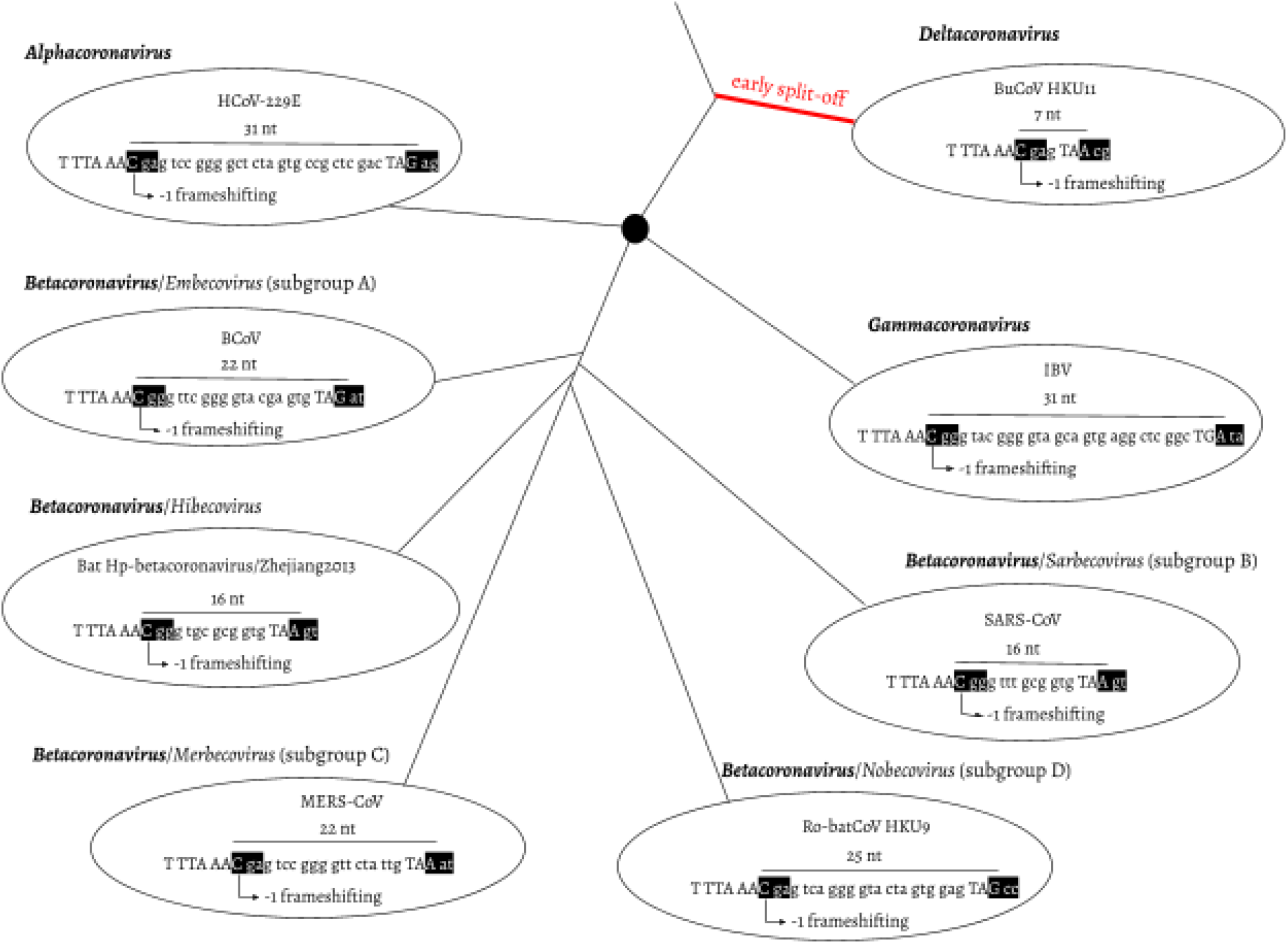
Overlap between ORF1a and b as a phylogenetic character. ORF1a frame 0 is shown as three consecutive nucleotides separated by spaces. The slippery site “TTT AAA C” and the ORF1a stop codon are shown in upper case letters. In ORF1b frame −1, two codons are shown in black boxes: the codon starting from the slippery site and the codon bypassing the ORF1a stop codon. *α-, β-*, and *γ-coronavirus* were plotted as splitting from one common node (black filled circle), with no phylogenetic order shown. HCoV-229E: Human coronavirus 229E, NC_002645.1. BCoV: Bovine coronavirus, NC_003045.1. Bat Hp-β-coronavirus/Zhejiang2013, NC_025217.1. MERS-COV, NC_019843.3. Ro-batCoV HKU9: Rousettus bat coronavirus HKU9, NC_009021.1. SARS-CoV, NC_004718.3. IBV: infectious bronchitis virus, NC_001451.1. BuCoV HKU11: Bulbul coronavirus HKU11, NC_011547.1.

**Fig. 2.**
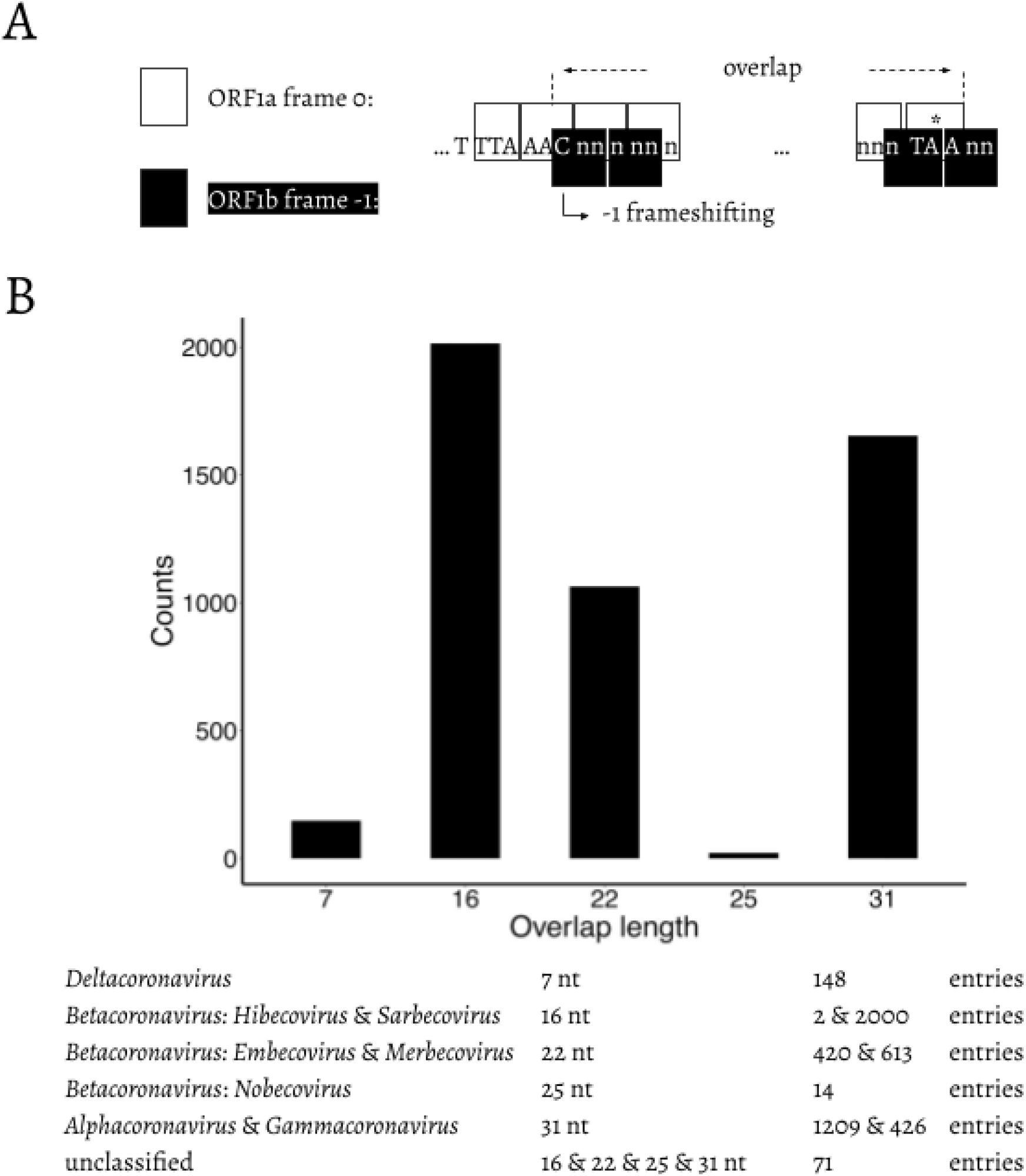
**A**. Schematic representation of the overlap between ORF1a and ORF1b. “TTT AAA C” is the slippery site. ORF1a frame 0 is shown in white boxes. “TTT AAA C” and the ORF1a stop codon are shown in upper case letters. ORF1b frame −1 is shown in black boxes. The −1 frameshifting starts from “C” of “TTT AAA C”, which is indicated at the bended arrow. * = the stop codon of ORF1a. **B**. Distribution of the length of the overlap in different genera/subgenera.

We then compared the first 15 amino acids of *ORF1b* in all 4,904 entries (Fig. 3). The amino acid sequences are highly conserved: positions 1 (R), 2 (V), 4 (G), 7 (S), 11-13 (ARL), and 15 (P) are almost invariable and highly redundant. Next, we compared underlying nucleotide sequences of the PFE region (Fig. 4). This suggests the following potential series of events. δ-coronavirus with 7 nt overlap most likely represents the ancestral state. Comparing coronaviruses with 7 nt (δ-coronavirus) and 31 nt (α- and γ-coronavirus) in the overlap, the stop codon to generate a 7 nt overlap is abolished at positions 5–7, through substitution events, which extends ORF1a to the next available stop codon at positions 38–40. This extension results in a new overlap with 31 nt in length (Fig. 4A). Comparing coronaviruses with 31 nt (α- and γ-coronavirus) and 25 nt (β-coronavirus: Nobecovirus) overlaps reveals a “GTA” insertion at positions 28–30. “TA” from the “GTA” together with the following “G” forms a new stop codon leading to a 31 → 25 nt shortening of the overlap. In a Nobecovirus with a 25 nt overlap, the 31 nt overlap stop codon (at positions 38–40) is still observable (Fig. 4B). Further comparison of coronaviruses with 31 nt (α- and γ-coronavirus) and 22 nt (β-coronavirus: Embecovirus and Merbecovirus) overlaps revealed a “GTA” insertion as well, but at positions 22–24. “TA” at positions 23–24 and the following “A” or “G” at position 25 constitute a new stop codon. In the 22 nt overlap, substitutions have been observed at the original stop codon (at positions 38–40) from 31 nt overlap coronaviruses; more specifically, “C” appears at position 39 (Fig. 4C). Finally, we compared coronaviruses with 31 and 16 nt length in the overlap. The same “GTA” insertion footprint was found, at positions 16–18 ahead of the two “GTA” insertions in 31 → 25 nt and 31 → 22 nt events. “TA” at positions 17–18 and the following “A” at position 19 form the stop codon in the 16 nt overlap coronaviruses. In addition, deletions at positions 13–15 were observed (Fig. 4D). These deletions are referred as “TCT”-like, since “TCT” are the dominant nucleotides observed at positions 13–15 in the 7 and 31 nt overlap coronaviruses. At positions 38–40, the ancestral stop codon in the 31 nt overlap coronaviruses can not be seen, since the nucleotide at position 39 is invariably represented by “T” (Fig. 4D). The variable position of the stop codon likely has an implication to the frameshift efficiency in these taxa as was shown by Bhatt et al. (Bhatt et al. 2021). These authors demonstrated that extension of the distance between the slippery heptamer and the stop codon of 0-frame decreases frameshifting frequency: an increase in the distance by 15 nucleotides, as is the case in α- and γ-coronaviruses (Fig. 4), decreases frameshifting by ~20%, while removal of the stop decreases it by half.

**Fig. 3.**
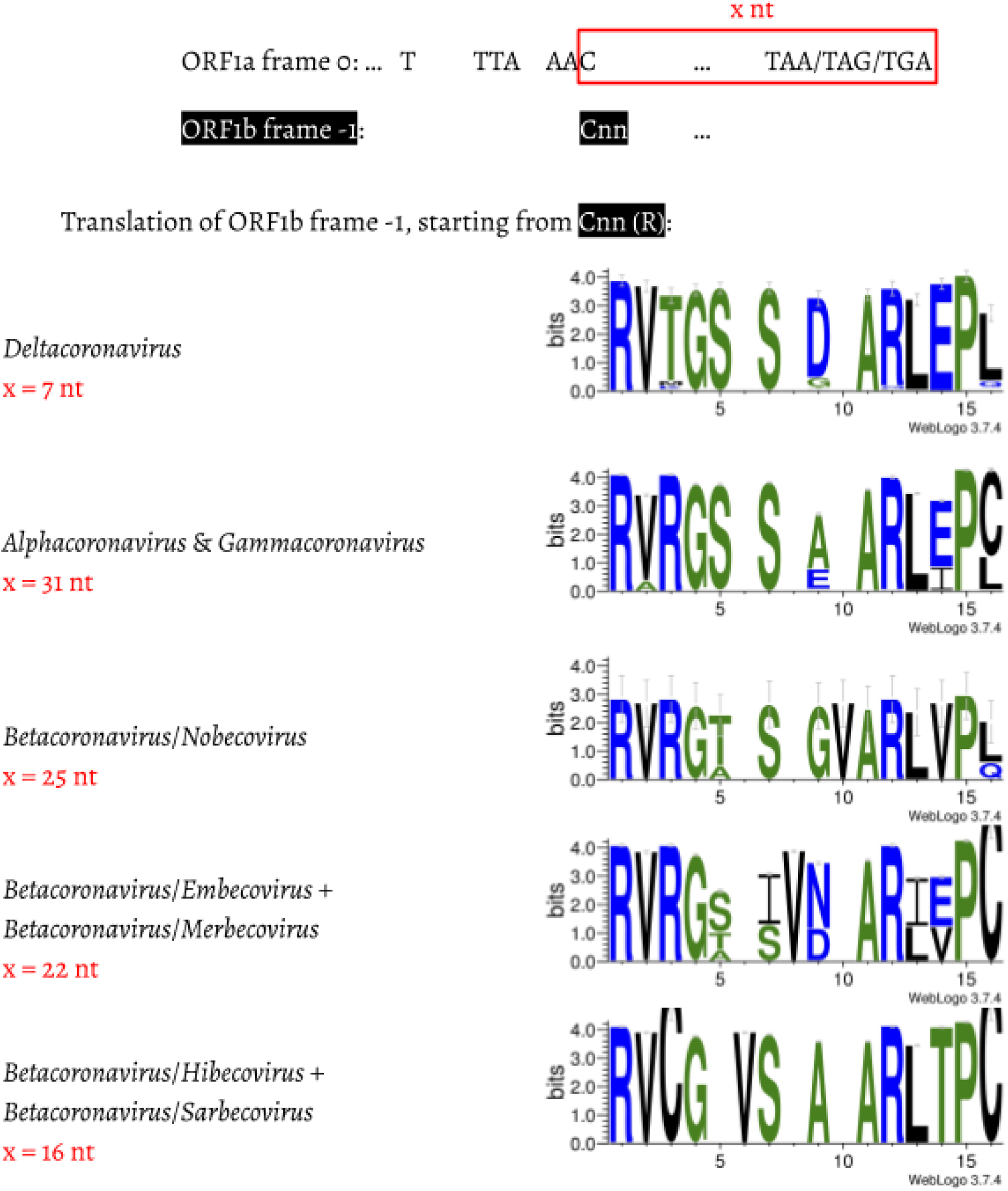
Amino acid alignment of the first 13–14 amino acids in coronaviruses with different lengths in the overlap region. For each genus/subgenus shown, all coronavirus entries belonging to this group are used to generate the consensus amino acid sequences. Gaps were left to make the amino acid sequences align.

**Fig. 4.**
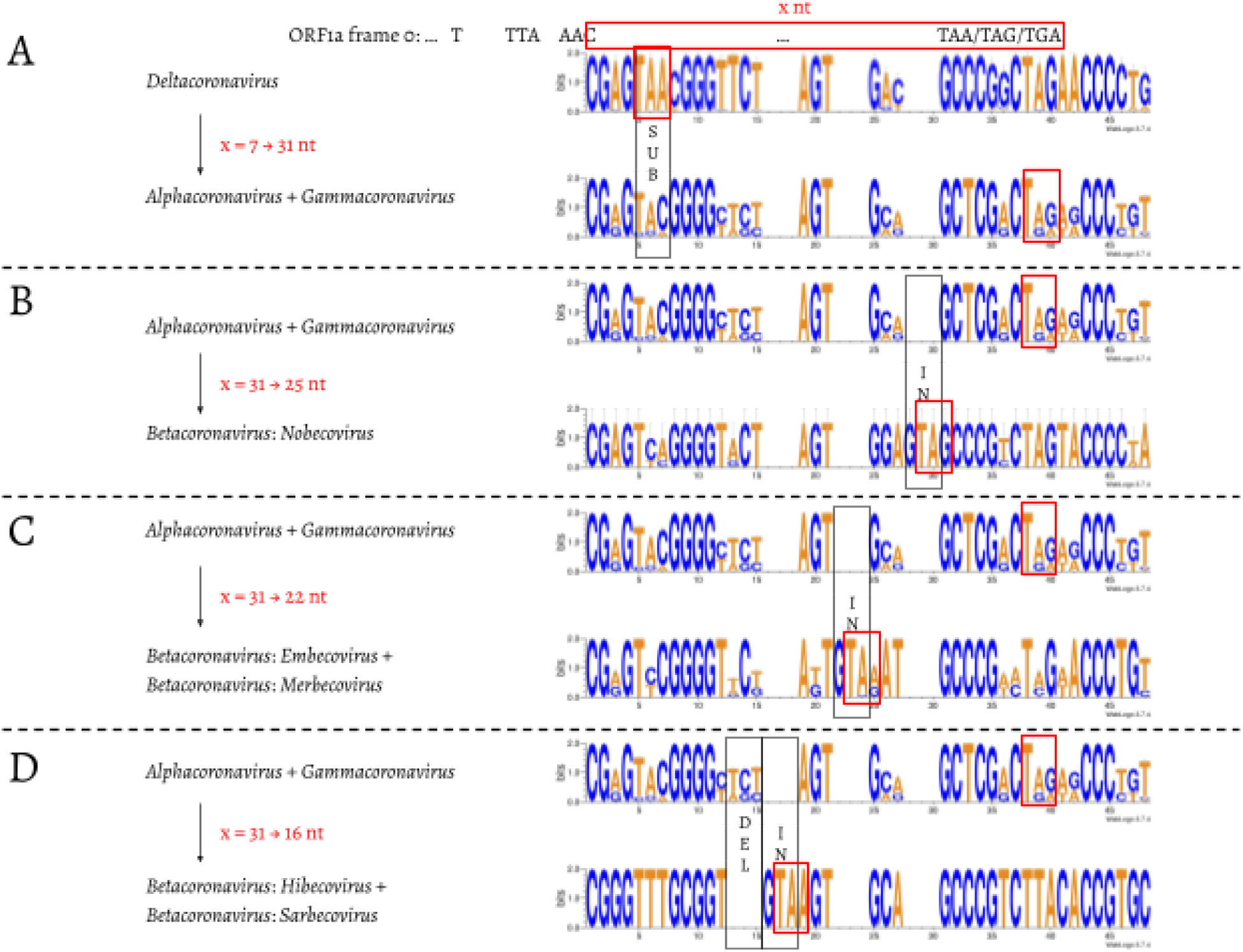
Nucleotide alignment of the overlap in coronaviruses with 7, 31, 25, 22, and 16 nt. The footprints of substitutions, insertions, and deletions are shown in black boxes, and labelled as “SUB”, “IN”, and “DEL”, respectively. The stop codon of ORF1a in each of the 7, 31, 25, 22, and 16 nt overlap coronaviruses is shown in a red box.

The abundance of SARS-CoV-2 sequencing data allows examining the substitution dynamics that may be present in population- and individual-level sequencing data. For population-level analysis we identified variants in the PFS region from >350,000 complete genome sequences available from GISAID (see Methods). However, because GISAID contains only assembled genomes, this data does not provide information about individual-level (intrasample) variation. For this we performed a detailed analysis of >55,000 samples generated with the COG-UK (Lythgoe et al. 2021) consortium (see (Maier et al. 2021) for analysis details). A summary of results from both analyses is shown in Table 1. There is little variation in the FSE region as the fraction of samples containing individual substitutions appears to be small (the two “Count” columns in Table 1). In addition, the vast majority of substitutions (30 out of 36 marked in Table 1) may be an artifact of RNA editing events from APOBEC (Chen and MacCarthy 2017) or ADAR (Bazak et al. 2014) enzymatic complexes. The remaining six substitutions (all transitions) are predominantly located in the loop regions of the predicted FSE secondary structure (Huston et al. 2021) and thus likely have no effect on the secondary structure.

**Table 1.** Allelic variants within the FSE region called from complete genomes (Population) and COG-UK (Individual) data. * = potential APOBEC-edited sites; + = potential ADAR-edited sites. Site numbering is in 0-based coordinates; † = out of 355,568 complete genomes; ‡ = out of 55,163 individual samples. Location of substitutions in a stem (^s^) or a loop (^L^) are based on structures predicted by Huston et al. (Huston et al. 2021).

| Site | Reference | Population |  | Individual |  |  |  |  |
| --- | --- | --- | --- | --- | --- | --- | --- | --- |
|  |  | Alternate | Count<sup>†</sup> | Alternate | Min AF | Max AF | Count<sup>‡</sup> | CoV |
| *13,425 | C | T | 131 | - | - | - | - | - |
| *13,429 | C | T | 178 | - | - | - | - | - |
| *13,430 | C | T | 57 | - | - | - | - | - |
| *13,431 | C | T | 18 | - | - | - | - | - |
| +13,432 | A | G | 23 | - | - | - | - | - |
| *13,434 | G | A | 33 | - | - | - | - | - |
| *13,435 | C | T | 48 | T | 0.116 | 0.971 | 14 | 0.403 |
| +13,437 | T | C | 7 | C | 0.985 | 0.988 | 5 | 0.001 |
| <sup>s</sup>13,440 | G | T | 6 | - | - | - | - | - |
| *13,445 | C | - | - | T | 0.068 | 0.970 | 25 | 0.239 |
| *13,447 | G | A | 6 | - | - | - | - | - |
| *13,451 | C | T | 25 | T | 0.941 | 0.977 | 19 | 0.009 |
| *13,457 | C | T | 251 | T | 0.052 | 0.963 | 19 | 0.842 |
| *13,458 | G | - | 246 | A | 0.069 | 0.970 | 6 | 0.669 |
| <sup>l</sup>13,458 | G | T | 246 | T | 0.080 | 0.976 | 6 | 0.442 |
| +13,481 | A | G | 9 | - | - | - | - | - |
| *13,486 | C | T | 336 | T | 0.055 | 0.965 | 7 | 1.012 |
| +13,487 | A | - | - | G | 0.901 | 0.949 | 12 | 0.017 |
| +13,497 | A | G | 6 | - | - | - | - | - |
| *13,498 | C | T | 16 | - | - | - | - | - |
| *13,500 | C | T | 53 | - | - | - | - | - |
| <sup>l</sup>13,504 | G | T | 6 | - | - | - | - | - |
| *13,505 | C | T | 7 | T | 0.887 | 0.917 | 5 | 0.015 |
| <sup>l</sup>13,511 | A | T | 13 | - | - | - | - | - |
| <sup>l</sup>13,512 | G | T | 13 | - | - | - | - | - |
| *13,513 | G | A | 21 | - | - | - | - | - |
| *13,514 | C | T | 41 | T | 0.065 | 0.889 | 6 | 0.622 |
| *13,516 | C | T | 711 | T | 0.101 | 0.840 | 49 | 0.260 |
| <sup>s</sup>13,525 | A | C | 46 | - | - | - | - | - |
| +13,526 | T | C | 14 | - | - | - | - | - |
| +13,532 | A | G | 6 | - | - | - | - | - |
| *13,535 | C | T | 3,999 | T | 0.215 | 0.841 | 23 | 0.223 |
| +13,541 | T | C | 6 | - | - | - | - | - |
| *13,547 | C | T | 59 | T | 0.067 | 0.898 | 8 | 0.373 |
| *13,550 | C | T | 31 | T | 0.878 | 0.921 | 11 | 0.017 |

Our results provide an alternative way to assess exceptional conservation of the PFE using publicly available sequence data highlighting the fact that the entire PFE region appears to be under strong purifying selection. These patterns are similar to observations obtained from deep mutational scanning where any alteration at the majority of PFE region sites results have deleterious effects on the frameshift efficiency (e.g., (Carmody et al. 2021)).

## Materials and Methods

### Coronavirus entries retrieval and filter

The 35,152 coronaviral entries in the NCBI taxonomy database were sorted by length, and only those larger than 14,945 nt were kept, leaving a total of 4,939 genomes. The slippery site and following overlap sequences were manually inspected, in case that the slippery site was incorrectly annotated. We further filtered out those entries if they contain no annotation information, or have gapped sequences in the overlap. By applying these filters, we finally had 4,904 coronavirus entries (Table S1), of which the overlap could be unambiguously determined.

### Amino acid alignment and nucleotide alignment of the overlap region

For all *δ-coronavirus* entries in Table S1, the first 13 amino acids of ORF1b were taken to generate a consensus sequence using WebLogo (Crooks et al. 2004). The same was done to *Alphacoronavirus* and *γ-coronavirus*. Within *β-coronavirus*, for *Nobecovirus, Embecovirus*, and *Merbecovirus*, the first 14 amino acids were used to build the consensus; for *Hibecovirus* and *Sarbecovirus*, the first 13 amino acids were used. In terms of the nucleotide sequence alignments, for each genus/subgenus, the nucleotide sequences used to generate the amino acids mentioned above were taken to make the nucleotide consensus sequence using WebLogo.

### Processing of GISAID data

Each genome was subjected to codon-aware alignment with the NCBI reference genome (accession number NC_045512) and then subdivided into ten regions based on CDS features: ORF1a (including nsp10), ORF1b (starting with nsp12), S, ORF3a, E, M, ORF6, ORF7a, ORF8, N, and ORF10. For each region, we scanned and discarded sequences containing too many ambiguous nucleotides to remove data with too many sequencing errors. Thresholds were 0.5% for the S gene, 0.1% for ORF1a and ORF1b genes, and 1% for all other genes. We mapped individual sequences to the NCBI reference genome (NC_045512) using a codon-aware extension to the Smith-Waterman algorithm implemented in HyPhy (Pond et al. 2005; Gianella et al. 2011) translated mapped sequence to amino-acids, and performed multiple protein sequence alignment with the auto settings function of MAFFT (version 7.453) (Katoh and Standley 2013). Codon sequences were next mapped onto the amino-acid alignment. Variants were called directly.

## Acknowledgements

The authors are grateful to Sergei Kosakovsky Pond for providing variant data from GISAID sequences. This work is funded by NIH Grants U41 HG006620 and NSF ABI Grant 1661497. The funders had no role in study design, data collection and analysis, decision to publish, or preparation of the manuscript.

